# Directed evolution of aerotolerance in sulfide-dependent thiazole synthases

**DOI:** 10.1101/2022.10.16.512417

**Authors:** Kristen Van Gelder, Edmar R. Oliveira-Filho, Jorge Donato García-García, You Hu, Steven D. Bruner, Andrew D. Hanson

## Abstract

Sulfide-dependent THI4 thiazole synthases could potentially be used to replace plant cysteine-dependent suicide THI4s, whose high protein turnover rates make thiamin synthesis exceptionally energy-expensive. However, sulfide-dependent THI4s are anaerobic or microoxic enzymes and hence unadapted to the aerobic conditions in plants; they are also slow enzymes (*k*_cat_ <1 h^-1^). To improve aerotolerance and activity, we applied continuous directed evolution under aerobic conditions in the yeast OrthoRep system to two sulfide-dependent bacterial THI4s. Six beneficial single mutations were identified, of which five lie in the active-site cleft predicted by structural modeling and two recapitulate features of naturally aerotolerant THI4s. That single mutations gave substantial improvements suggests that further advance under selection will be possible by stacking mutations. This proof-of-concept study established that the performance of sulfide-dependent THI4s in aerobic conditions is evolvable and, more generally, that yeast OrthoRep provides a plant-like bridge to adapt nonplant enzymes to work better in plants.

## INTRODUCTION

Plants and fungi synthesize the thiazole moiety of thiamin using a suicide THI4 thiazole synthase (EC 2.4.2.60) in which an active-site Cys residue donates the required sulfur atom; this irreversibly inactivates the THI4 after just one reaction cycle (**Figure 1**).^1,2^ The energy cost of replacing an inactivated plant THI4 is equivalent to a biomass yield penalty of up to 2-4%.^3^ Exchanging a Cys-dependent suicide THI4 for a catalytic THI4 (EC 2.4.2.59) that mediates multiple reaction cycles using sulfide as sulfur donor (**Figure 1**)^2,4^ could therefore potentially increase biomass yield, making this a rational synthetic biology strategy for crop improvement.^3,5^ However, there are obstacles to doing this. Sulfide-dependent THI4s, which have an Fe(II) cofactor,^4^ come from organisms whose environments are anoxic or deeply hypoxic, sulfide-rich, and in some cases, hot (≥60°C)^4,6–8^ and so are ill-adapted to plants.^8^ One challenge is thus to modify sulfide-dependent THI4s to work well in air (*vs*. <1% oxygen), low-μM sulfide^9^ (*vs*. high-μM to mM^8^), and mild temperatures. Another challenge is that sulfide-dependent THI4s are slow enzymes: *Methanococcus jannaschii* THI4 mediated only two catalytic cycles *in vitro* in 3.5 h^4^ (i.e., *k_cat_* <1 h^-1^, a catalytic turnover rate so low that it precludes biochemical characterization). Similarly, while sulfide-dependent THI4s complemented an *Escherichia coli* thiazole auxotroph in aerobic conditions,^8^ the THI4 activity per unit protein needed for this was very low since the THI4s were ~30% of soluble protein^9^ and even suicide THI4s complement *E. coli* if expressed at such levels.^2,10^

**Figure 1.**
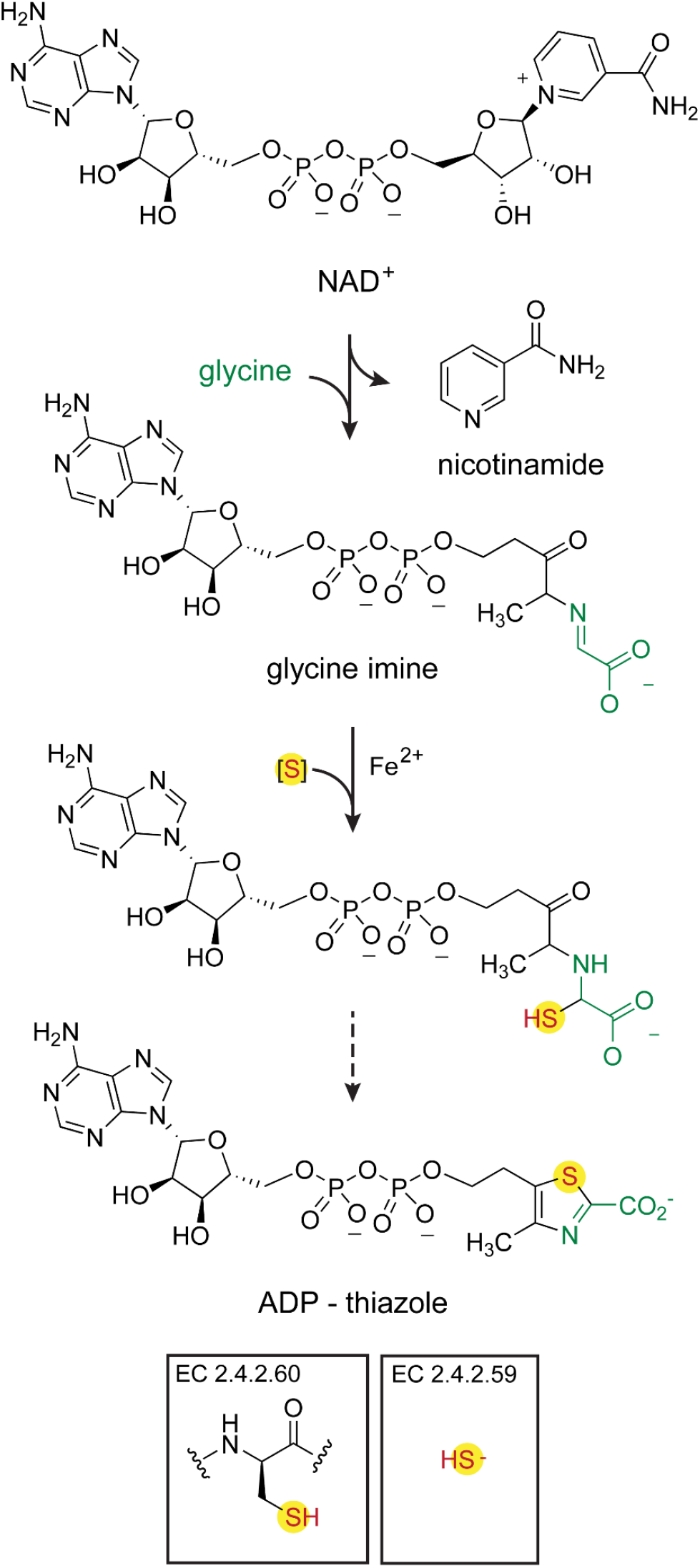
Biosynthesis of adenylated carboxythiazole (ADP-thiazole) by cysteine-dependent and sulfide-dependent THI4s. ADP-thiazole is formed from NAD^+^, glycine, and a sulfur atom. Cysteine-dependent THI4s take the sulfur atom from an active-site cysteine residue, leaving a dehydroalanine residue and causing inactivation. Sulfide-dependent THI4s use sulfide as sulfur donor.

This study aimed to adapt sulfide-dependent THI4s to aerobic conditions using continuous directed evolution (CDE), which has unmatched capacity for mutational depth and scale.^11,12^ In CDE, the enzyme gene is hypermutated *in vivo*, the enzyme’s activity is coupled to host cell growth, and improved variants are obtained by selecting for faster growth.^12^ We chose the yeast *(Saccharomyces cerevisiae)* OrthoRep CDE system because: (i) it durably mutates a target gene at ~10^5^-fold the natural rate;^13,14^ (ii) sulfide-dependent prokaryote THI4s complement a yeast thiazole auxotroph;^8^ and (iii) yeast is a closer facsimile of plants than *Escherichia coli*,^15^ the main alternative CDE platform.^12^ To avoid overtaxing the initially low activity of the target THI4s, we supported their function by using a host strain *(met15Δ*) with a high internal sulfide level.^16^ Since cytosolic conditions in aerobic yeast, as in plants, differ from those in anaerobic bacteria with respect to, e.g., NADH/NAD^+^ ratio^17–19^ and protein folding and degradation systems,^15^ selection in yeast might improve adaptation to such factors as well as to oxygen. Structure modeling and comparative genomics were applied to distinguish these possibilities.

## RESULTS AND DISCUSSION

### Target choice and pre-screening

We chose three bacterial THI4s that have detectable activity under aerobic, high-sulfide conditions, as shown by complementation of an *E. coli*Δ*thiG* thiazole auxotroph.^8^ These were: MhTHI4 from *Mucinivorans hirudinis* (leech gut mesophile); SfTHI4 from *Saccharicrinis fermentans* (marine mud mesophile); and TaTHI4 from *Thermovibrio ammonificans* (deep-sea vent thermophile). As OrthoRep expresses target genes at relatively low levels,^12^ we pre-screened MhTHI4 and SfTHI4 for ability to complement a yeast *met15*Δ *thi4*Δ strain when more highly expressed from a CEN6/ARS4 plasmid, as done for TaTHI4.^8^ (If complementation fails this pre-screen it will fail in OrthoRep.) The THI4s were tested plus or minus yeast THI4’s 28 N-terminal residues, a putative mitochondrial targeting peptide. All three THI4s complemented and so went forward to OrthoRep. The yeast N-terminus had little effect on complementation (**Figure S1**) and was not added in further experiments.

### CDE in OrthoRep

We transformed a *met15*Δ *thi4*Δ strain with the nuclear ArEc-TDH3 plasmid harboring the error-prone polymerase TP-DNAP1_611, then introduced plasmids p1 and p2 by protoplast fusion.^15^ p1 harbors the *THI4* target gene that the error-prone polymerase hypermutates (**Figure 2A**). We ran three independent clones of each THI4 through three selection schemes:^15^ 1, serial passages on thiamin-free medium; 2, initial passages on medium with limiting thiamin, then transfer to thiamin-free medium; and 3, initial passages on medium with luxury thiamin (to build a mutant library), then transfer to scheme 2 (**Figure 2B–D**). Six MhTHI4 and three SfTHI4 populations showed growth in one or more schemes; growth improved with passaging until it neared that of a control whose p1 harbored yeast *THI4* (**Figure 2B–D** and **Figure S2**). As this control marks the upper limit of the selection window, we ended campaigns at this point and sequenced individual clones from each population. No TaTHI4 population showed any growth, possibly because this thermophilic THI4 had too little activity at 30°C.

**Figure 2.**
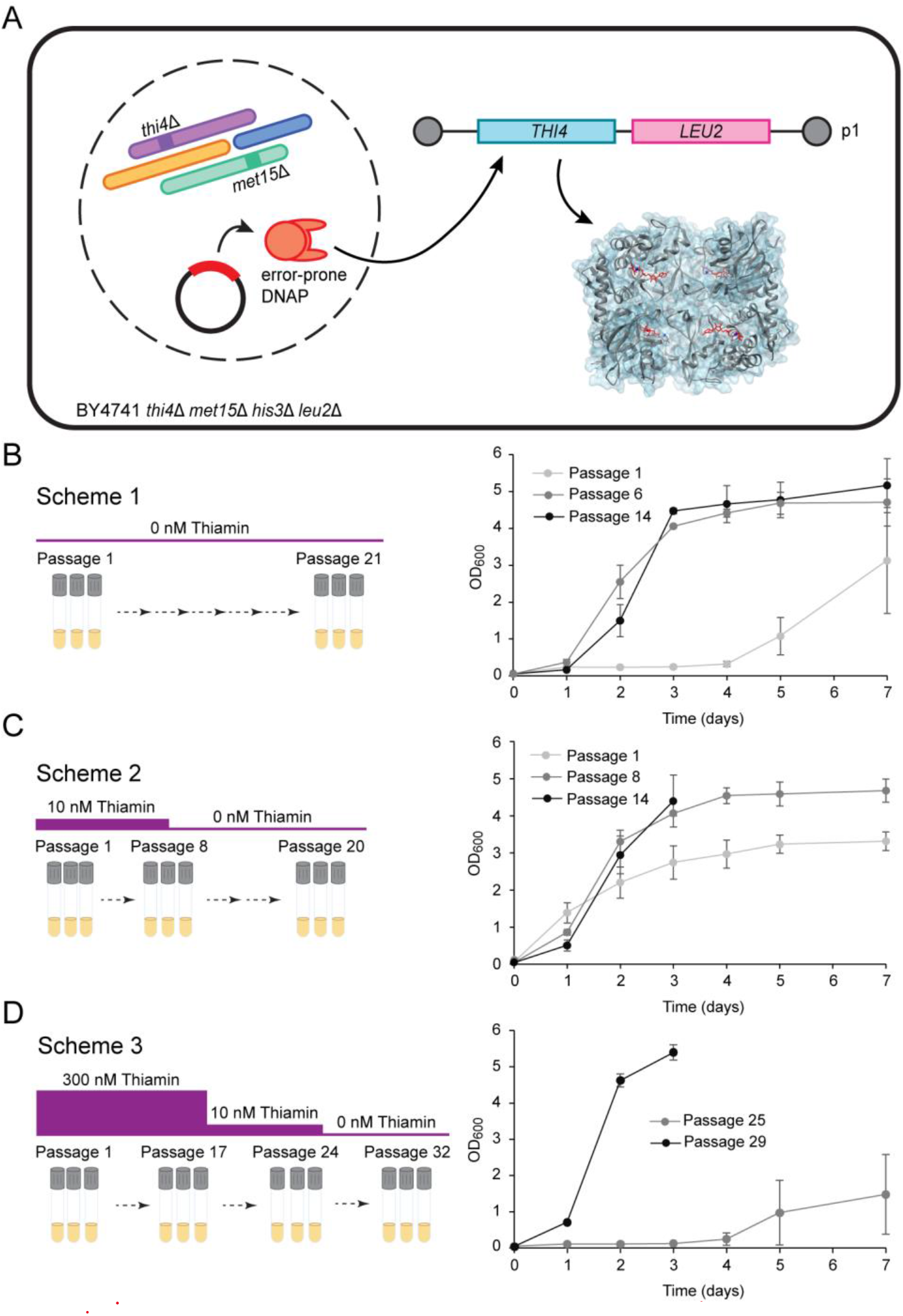
Continuous directed evolution of MhTHI4 and SfTHI4 in OrthoRep. (A) Overview of Ortho-Rep. An error-prone version of the terminal protein-primed DNA polymerase (TP-DNAP1) of the p1 plasmid is transferred to a nuclear plasmid with a *HIS3* selection marker in strain BY4741 *thi4△ met15△ his3△ leu2△ ura3△*. TP-DNAP1 on p1 is replaced by the *MhTHI4* or *SfTHI4* gene and a *LEU2* marker. The error-prone polymerase replicates p1, mutating the *THI4*. (B) Selection scheme 1. MhTHI4 and SfTHI4 were evolved without thiamin for up to 21 passages. Growth (OD_600_) of 3-mL cultures was monitored and is shown for passages 1, 6, and 14. (C) Selection scheme 2. MhTHI4 and SfTHI4 were evolved for eight passages in 10 nM thiamin, then for 12 passages without thiamin. Growth is shown for passages 1 and 8 (10 nM thiamin) and passage 14 (no thiamin). (D) Selection scheme 3. MhTHI4 and SfTHI4 were grown for 17 passages in 300 nM thiamin, in 10 nM thiamin for seven passages, and without thiamin for eight passages. Growth is shown for passages 25 and 29 (no thiamin). All data are means ± s.e.m. of 2-3 replicates.

### CDE outcomes

Sixteen variants carrying one or two nonsynonymous ORF mutations were recovered, mostly from MhTHI4 (**Figure 3A**), as well as variants with synonymous mutations and promoter mutations (**Figure S3**). ORF mutations were 75% T>C, 15% A>G, and 10% G>A, which is similar to those reported.^13^ Four single MhTHI4 mutations recurred in independent populations: V124A in three, and V28A, Y122C, and V138A in two each. Likewise, the SfTHI4 mutation D168G recurred in three populations. Such convergent evolution of mutations implies fitness benefits. To confirm that benefits were due to mutations in the *THI4* ORF and not elsewhere in p1 or the yeast genome, each mutant ORF was recloned into p1, inserted into fresh cells, and growth in thiamin-free medium was measured for independent clones. Wildtype MhTHI4 or SfTHI4 and yeast THI4 served as benchmarks to gauge improvement (**Figure 3B** and **Figure S4**). Taking MhTH4 and SfTHI4 together, six mutations were beneficial, five neutral, and four deleterious, i.e., acted alone or cancelled a beneficial mutation (**Figure 3B**). Consistent with expectation, all convergent mutations were beneficial. The deleterious mutations probably arose after beneficial ones had partly swept the population (**Figure 3A**) and survived because the cells harboring them were cross-fed thiamin by thiamin-producers in the rest of the population.^15^ Because sulfide-dependent THI4s are labile and have extremely low *k*_cat_ values,^4,8^ the biochemical characteristics of wildtype and evolved enzymes cannot be compared using *in vitro* assays. We therefore turned to comparative genomics and structure modeling to interpret the observed mutations.

**Figure 3.**
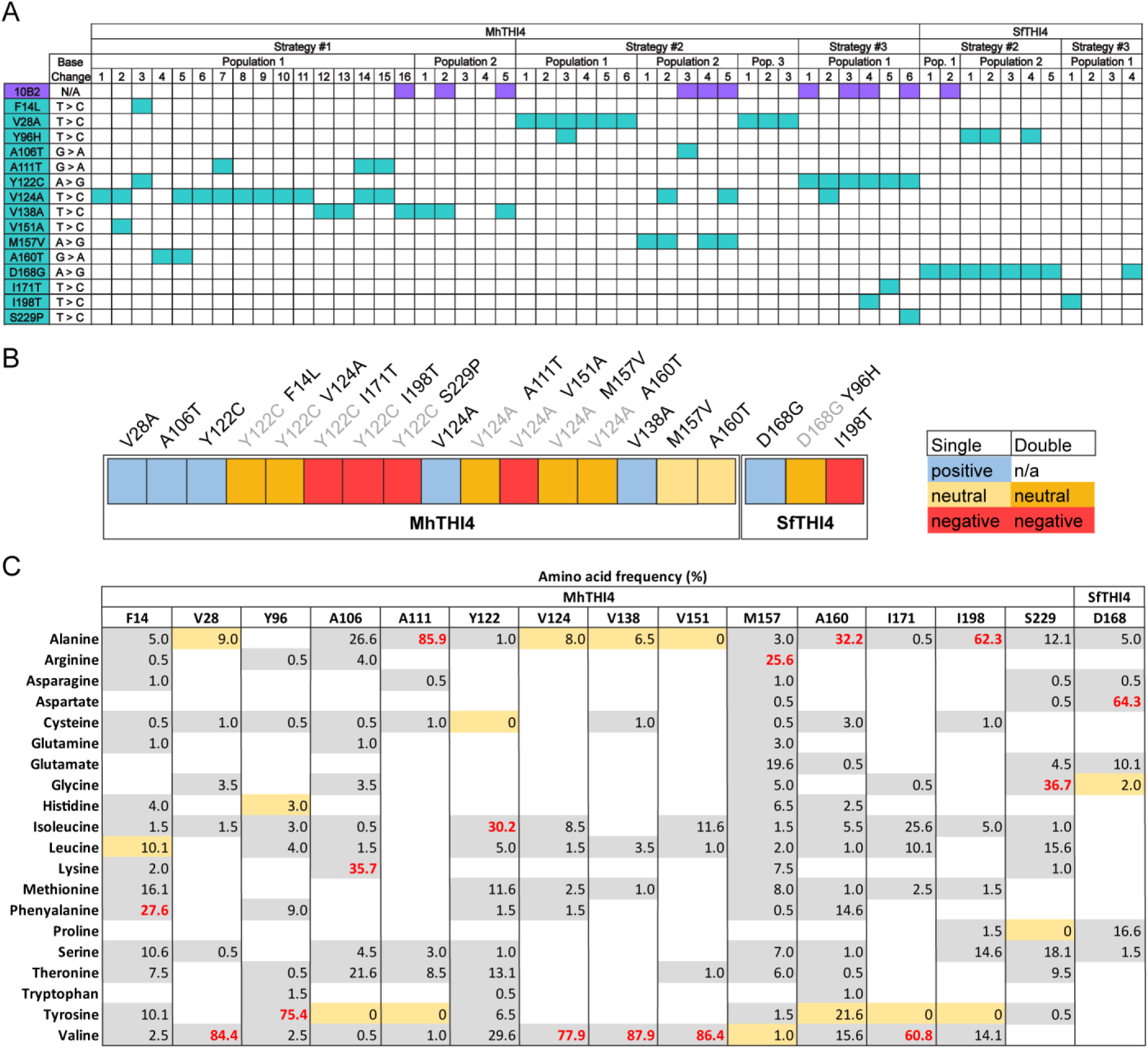
Mutations in six evolved populations of MhTHI4 and three of SfTHI4. (A) Promoter (10B2) and ORF mutations. V124A and D168G each occurred in three populations; V28A, Y96H, Y122C, and V138A each occurred in two populations. (B) Classification of mutations based on growth relative to that supported by wildtype MhTHI4 or SfTHI4 and yeast THI4 as benchmarks. Beneficial (positive) mutations grew approximately as well as the yeast benchmark. Deleterious (negative) mutations prevented growth when alone or when combined with a beneficial mutation. Neutral mutations had no measurable effect. Double mutants are classified according to how the second mutation (black) affected the first positive mutation (gray). (C) Natural variation in residues at mutation sites. Numbers are frequencies (%) of naturally occurring amino acids at each site among 199 diverse sulfide-dependent THI4s;^8^ the most frequent amino acid is in red. White spaces indicate 0% frequency. The observed mutations are highlighted in beige.

### Mutations in relation to natural variation

We assessed natural variation among residues at the mutated positions using a set of 199 diverse sulfide-dependent THI4s^8^ (**Figure 3C**). Of the six beneficial mutations, all except Y122C and D168G are conservative or semi-conservative, and all except Y122C occur naturally at frequencies of 2-22%. The five neutral mutations are also conservative and occur naturally. In contrast, two of the deleterious mutations are nonconservative and none are natural. The outcomes of the CDE campaigns are thus broadly similar to the outcomes of natural selection.

### Structure modeling

The mutations’ positions were mapped onto the modeled structure of MhTHI4 (which is very similar to that of SfTHI4) (**Figure 4A**). The five convergent, beneficial mutations (V28A, Y122C, V124A, V138A, and D168G) are in the active-site cleft, near the adenine ring of the bound intermediate. The sixth beneficial mutation, A106T, is in the dimer interface. Two of the four deleterious mutations (I171T and V151A) are in or near the active-site and two (I198T and S229P) are on the protein surface. The neutral mutations are all in or near the dimer interface or on the surface. Interestingly, beneficial mutations V28A, V124A, and V138A line a hydrophobic pocket via which oxygen might access the Fe(II) center. Replacing any of them with a smaller Ala residue could allow closer packing that impedes oxygen access and thus lessens damage to the catalytic metal. Another interesting possibility is that the beneficial mutations near the bound intermediate reduce discrimination between the NAD^+^ substrate and NADH, which might compete with NAD^+^ for binding at the active-site. Such discrimination may be less necessary when MhTHI4 and SfTHI4 are expressed in yeast instead of their anaerobic native hosts, whose cytosolic NADH/NAD^+^ ratios are likely at least tenfold that in yeast.^17,18^

**Figure 4.**
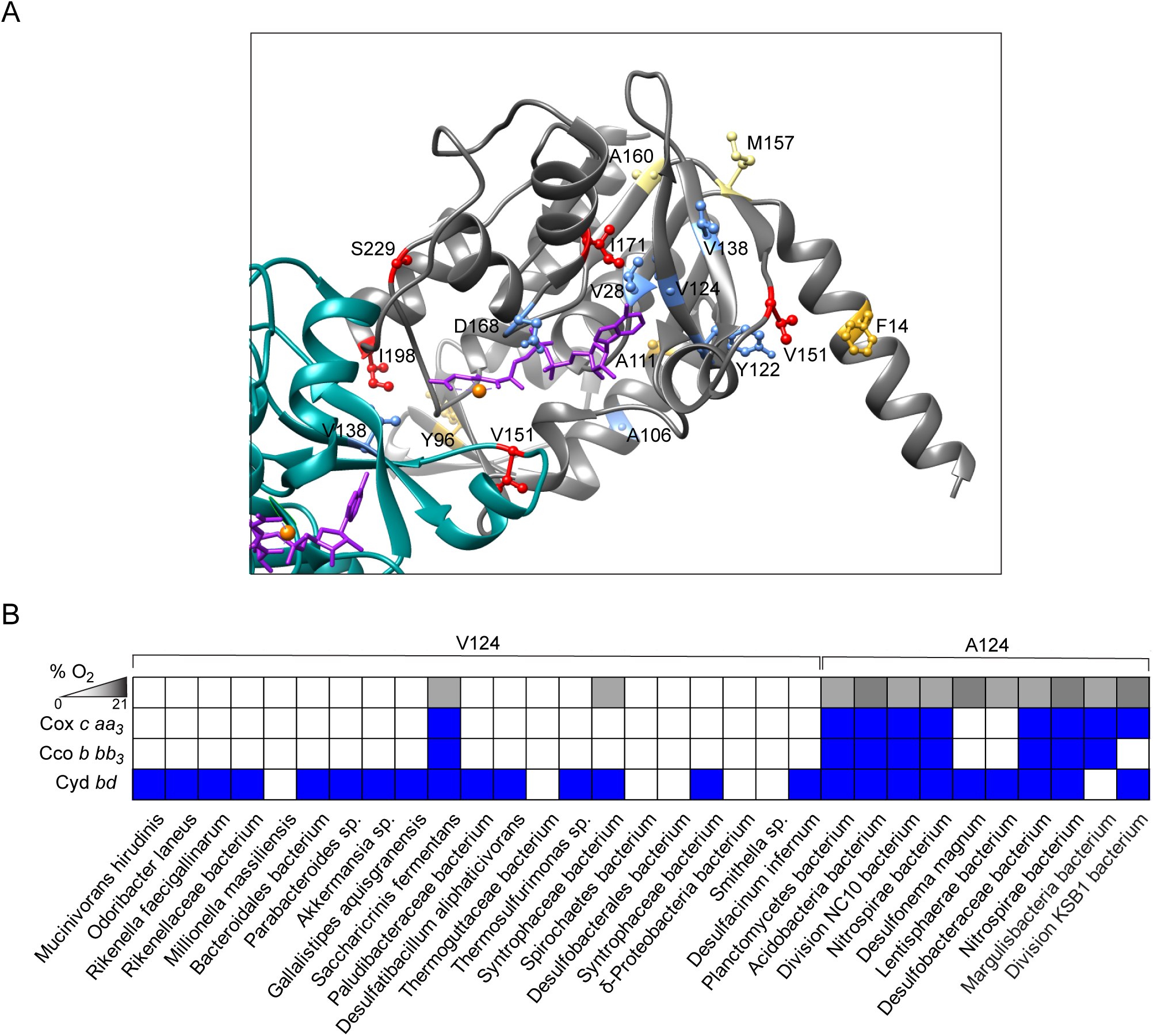
The positions of beneficial, neutral, and deleterious mutations within the modeled MhTHI4 structure, and the association of the natural V124A variant with oxygen-tolerance. (A) The observed mutations mapped onto the structure of MhTHI4, modeled from the crystal structure of *Thermovibrio ammonificans* THI4 (PDB: 7RK0). Two subunits of the octamer are shown (subunit A is gray, subunit B is cyan) with mutations indicated in blue (beneficial), gold (neutral, double), khaki (neutral, single) and red (deleterious). The bound substrate is indicated in purple and the metal ion in orange. B) The occurrence of V124 and A124 variants in natural THI4 sequences in relation to the oxygen adapt-ation of the host organism, based on habitat data and on the types of cytochrome encoded by the genome, i.e., *aa*_3_-type and *cbb*_3_-type cytochrome *c* oxidases and cytochrome *bd*, which are adapted to operate at progressively lower oxygen levels.^20^ Habitat oxygen is shown schematically on a gray scale, from zero (white) to air-level (black). See **Figure S5** for a larger panel of oxygen metabolism genes and more detailed information.

### Comparative genomics of beneficial mutations

Comparative genomics suggests rationales for three of the beneficial mutations. The first is V124A, a fairly common natural variant (**Figure 3C**) that comparative genomics implicates in oxygen-tolerance, based on the presence or absence of genes for oxygen-metabolizing enzymes (**Figure S5**). Most simply, if A at position 124 confers more oxygen tolerance than V, then V124 should be prevalent in low-oxygen organisms and A124 in high-oxygen ones. This is observed (**Figure 4B** and **Figure S5**). Most A124-THI4s are from organisms of hypoxic or microoxic habitats whose genomes encode *aa*_3_-type and *cbb*_3_-type cytochrome *c* oxidases, which are adapted to high and low oxygen levels, respectively.^20^ Conversely, most V124-THI4s are from anaerobes that lack cytochrome *c* oxidases and whose sole oxygen-dependent terminal oxidase is cytochrome *bd*, which is adapted to ultralow oxygen levels and may serve to scavenge trace oxygen.^20^

The non-natural MhTHI4 Y122C mutation might also improve oxygen tolerance. Changing Y122 to C creates, with C121, a CC motif near the Fe(II) center that mimics one in the aerobic suicide THI4 of *Ustilago trichophora* and other fungi. As Cys residues can provide sacrificial thiol groups to protect critical sites from oxidants,^21^ the Y122C mutation might shield the Fe(II) center from oxidation. An alternative rationale is that a small hydrophobic residue is preferred at position 122 (**Figure 3C**), and Cys is the only such residue accessible from the Y122 codon (TAC) by an OrthoRep point mutation.^13^

The SfTHI4 D168G mutation could confer a benefit unrelated to oxygen. D168 is ~10 Å from the Fe(II) center and is followed by C. DC motifs can be hotspots for spontaneous isomerization of aspartate to isoaspartate, which alters the peptide backbone structure and can reduce enzyme activity.^22^ Like most other organisms, *S. fermentans* (the source of SfTHI4) has the protein isoaspartyl methyltransferase (PIMT) that repairs isoaspartyl residues; yeast does not.^23^ Thus, despite being a radical mutation, D168G could benefit yeast, but *only* yeast, by reducing isomerization damage that yeast cannot repair.

### Performance of beneficial mutations in *E. coli*

Consistent with the above possibility, in complementation tests in *E. coli* (which has PIMT) the D168G variant was far less active than wildtype SfTHI4 (**Figure S6**), i.e., the D168G mutation was deleterious rather than beneficial in a host that repairs isoaspartyl residues. The non-natural Y122C mutation was moderately deleterious in *E. coli* whereas the V124A mutation was neutral (**Figure S6**). That the V124A mutant did not outperform the wildtype could be because shake cultures of *E. coli*are more severely hypoxic than those of yeast,^24^ i.e., improved oxygen tolerance may have conferred no growth advantage in *E. coli* in our culture conditions.

### Deleterious mutations

Because they are common and *a priori* unlikely to specifically affect oxygen tolerance, detrimental mutations were not examined in depth. Two nevertheless stood out: I198T and V151A (**Figures 3B** and **4A**). The I198T mutation abolished activity when present alone in SfTHI4 and when combined with the beneficial Y122C mutation in MhTHI4. As I198 is a surface residue that interacts with the adjacent monomer’s N-terminus, replacement by Thr could impact dimerization. The V151A mutation in MhTHI4 abrogated the benefit of V124A and abolished activity. Because V151 is next to an Fe(II)-liganding Asp^8^, its replacement by Ala could disrupt metalation.

## CONCLUSIONS

Our evolution campaigns delivered single-mutant THI4s that drove growth rates similar to yeast native THI4, so that only the sequence space within one mutational step of the wildtype sequence could be explored. There is thus much space left to explore to improve activity in aerobic, plant-like conditions. The THI4s evolved so far are *a priori* likely to be improvable. First, our improved THI4s at best equalled the performance of yeast THI4. This is a low bar: as yeast THI4 is a suicide enzyme^1^ with a half-life of ~9 h,^25^ improved THI4s need make *only a few turnovers per day* to match its activity. The median enzyme turnover number (*k*_cat_) is 10^5^-fold more than this.^26^ Second, we have not yet selected for activity in low-sulfide conditions, or for Co(II) vs. Fe(II) as cofactor, which could reduce oxygen-sensitivity.^4,8^

Beyond the quest for oxygen-tolerance *perse*, this study sought proof-of-concept for OrthoRep in two new applications: (i) as a plant-like bridge to evolve microbial enzymes to function better in plants, for which there is an unmet need;^15,27^ and (ii) as a platform to increase the number of catalytic cycles that enzymes mediate *in vivo* before being replaced (catalytic-cycles-until-replacement, CCR^28^). The effectiveness of OrthoRep demonstrated here supports its use in both these applications.

## METHODS

### Plasmid construction

THI4 cloning in *E. coli* strain DH10B/TOP10 was as previously described.^14,15^ Bacterial THI4s were codon-optimized for yeast and synthesized by GenScript (Piscataway, NJ); the recoded sequences are given in **Table S1**. Native yeast THI4 was not recoded. The recoded bacterial THI4s were used with or without addition of the N-terminal 28 codons of yeast THI4 plus a GG linker (encoding MSATSTATSTSASQLHLNSTPVTHCLSDGG). Plasmids used in this study are listed in **Table S2**. Primers were purchased from Eurofins Genomics (Louisville, KY) and are listed in **Table S3**. Enzymes for PCR and cloning were obtained from Thermo Fisher Scientific (Miami, FL).

### Yeast strains and media

Strains are listed in **Table S4**. Yeast was grown in YPD or selection media as described.^15^ Selection media were synthetic complete (SC) minus the amino acids used as selection markers, e.g., yeast GA-Y319 containing *p1_THI4* was grown in SC-Leu and yeast BY4741 *thi4Δ* containing *p1_THI4* and the error-prone TP-DNAP was grown in SC-His-Leu-Trp. Before cloning THI4s in OrthoRep, complementation in BY4741 *thi4Δ* was confirmed by expression in ArEC-TDH3^15^.

### Yeast transformation and protoplast fusion

Transformations were performed as previously^15^ with minor modifications. ScaI-digested GR-306MP harboring *ScTHI4, MhTHI4*, or *SfTHI4* was added to GA-Y319 competent cells with 10 μL of 11 mg/mL ssDNA and 600 μL of an 8:1:1 50% PEG:10× TE:1 M lithium acetate mixture and incubated for 45 min at 30°C. Following an additional 20 min incubation at 42°C, the DNA-cell mixture was centrifuged (5 min, 700*g*), resuspended in 1 mL of sterile MilliQ water and plated on SC-Leu. Plates were incubated at 30°C for up to 5 days and single colonies were selected for gDNA isolation and sequencing to confirm integration of the *THI4* gene and the leucine selection marker in the p1 plasmid.^15^ Sequence-verified *p1_THI4* in GA-Y319 was used as the donor strain for protoplast fusion and BY4741 *thi4Δ* as recipient. Protoplast fusions were carried out as before.^15^

### Evolution campaigns

Campaigns used the three schemes described previously^15^ and shown in **Figure 2B–D**. Two MhTHI4 populations were evolved by scheme 1, three MhTHI4 populations and two SfTHI4 populations were evolved by scheme 2, and one population each of MhThi4 and SfTHI4 was evolved by scheme 3. Cells were washed five times with thiamin-free medium when transitioning from thiamin-containing to thiamin-free medium. Cultures were started at OD_600_ = 0.05. Cultures were analyzed for mutations by sequencing amplicons containing the THI4 ORF and promoter. Mutant THI4s were recloned in p1, introduced into fresh cells, and validated as described^15^ and outlined in the text.

### Comparative genomics

Distributions of V124-THI4s and A124-THI4s and oxygen metabolism genes were analyzed using GenBank complete genomes and metagenome assembled genomes and Blastp. The query sequences used for oxygen metabolism genes are given in the legend of **Figure S5**.

### Structure modeling

The MhTHI4 model was generated using Swiss-Model (Expasy) with the TaTHI4 structure as template (PDB: 7RK0). The homology model had a global model quality estimate of 0.86.^29^ Structure graphics were made using Chimera 1.15.^30^

## Author Contributions

^1^A.D.H. devised the study. K.V.G., J.D.G.-G., and E.R.O.-F. ran experiments; K.V.G., Y.H., and S.D.B. built structure models. All authors analyzed data. A.D.H. and K.V.G. wrote the manuscript.

## ACKNOWLEDGEMENTS

We thank M.A. Wilson and N. Smith for helpful discussions. This work was supported primarily by the U.S. Department of Energy, Office of Science, Basic Energy Sciences under Award DE-SC0020153 (to A.D.H.), and by USDA National Institute of Food and Agriculture Hatch project FLA-HOS-005796, and an Endowment from the C.V. Griffin, Sr. Foundation.

